# Selection constrains phenotypic evolution in a functionally important plant trait

**DOI:** 10.1101/015172

**Authors:** Christopher D. Muir

## Abstract

A long-standing idea is that the macroevolutionary adaptive landscape – a ‘map’ of phenotype to fitness – constrains evolution because certain phenotypes are fit, while others are universally unfit. Such constraints should be evident in traits that, across many species, cluster around particular modal values, with few intermediates between modes. Here, I compile a new global database of 599 species from 94 plant families showing that stomatal ratio, an important functional trait affecting photosynthesis, is multimodal, hinting at distinct peaks in the adaptive landscape. The dataset confirms that most plants have all their stomata on the lower leaf surface (hypostomy), but shows for the first time that species with roughly half their stomata on each leaf surface (amphistomy) form a distinct mode in the trait distribution. Based on a new evolutionary process model, this multimodal pattern is unlikely without constraint. Further, multimodality has evolved repeatedly across disparate families, evincing long-term constraint on the adaptive landscape. A simple cost-benefit model of stomatal ratio demonstrates that selection alone is sufficient to generate an adaptive landscape with multiple peaks. Finally, phylogenetic comparative methods indicate that life history evolution drives shifts between peaks. This implies that the adaptive benefit conferred by amphistomy – increased photosynthesis – is most important in plants with fast life histories, challenging existing ideas that amphistomy is an adaptation to thick leaves and open habitats. I conclude that peaks in the adaptive landscape have been constrained by selection over much of land plant evolution, leading to predictable, repeatable patterns of evolution.

---

The topography of the macroevolutionary adaptive landscape is thought to shape the broad patterns of life’s diversity [1]. Adaptive landscapes with multiple peaks are manifest in convergent evolution of similar phenotypes across independent evolutionary lineages. In such cases, surveys across species should reveal a multimodal trait distribution in which the modes point to the underlying peaks in the landscape. Multimodality has been observed frequently among plants and animals, including traits such as self-incompatibility [2], the precocial-altricial spectrum [3], pollination syndromes [4], ecomorphology in *Anolis* [5], and plant height [6]. That such disparate classes of traits show broadly similar patterns suggests that divergence on a multi-peaked adaptive landscape may be a general feature of macroevolution. However, we rarely know whether multimodality reflects constraints imposed by the underlying adaptive landscape and not some other constraint on phenotypic evolution.

In particular, certain phenotypes may be common not because they are more fit, but rather because they are genetically, developmentally, or functionally accessible. Conversely, rare phenotypes might be inaccessible. I use the definitions given by Arnold [7]: genetic constraints are limitations set by the “pattern of genetic variation and covariation for a set of traits”; developmental constraints are limitations on “possible developmental states”; and functional constraints are imposed by “time, energy, or the laws of physics”. Arnold contrasts these with selective constraints determined by the adaptive landscape. There are examples of genetic [8], developmental [9], and functional [10] constraints on phenotypic evolution acting in nature, meaning that we cannot assume selection alone shapes trait evolution. Compelling evidence from cross species comparisons that selection constrains phenotypic evolution requires showing that phenotypic evolution is constrained, that selection is sufficient to explain the inferred constraint, and that nonselective constraints are inconsistent with these observations.

Here, I evaluate evidence for selective constraints on a functionally important plant trait, stomatal ratio, using comparative methods and theory. Stomatal ratio, defined as the ratio of upper to lower stomatal density, impacts how plants ‘eat’ (i.e. assimilate CO_2_ from the atmosphere via photosynthesis). Physiological experiments and biophysical theory demonstrate that amphistomatous leaves, those that have equal stomatal densities on both upper and lower surfaces, maximize photosynthetic rate by minimizing the distance between substomatal cavities and chloroplasts, facilitating rapid CO_2_ diffusion [11, 12, 13, 14]. Hence, nearly all plants should be amphistomatous to maximize photosynthesis, yet paradoxically up to 90% of plant species in some communities are hypostomatous [15, 16, 17, 18], meaning that most stomata are on the lower surface. In rare cases, most stomata are on the upper surface (hyperstomy). I use upper and lower rather than abaxial and adaxial, because the former applies to ‘upside-down’ (i.e. resupinate) leaves. Stomatal ratio is a quantitative metric that describes continuous variation between hypo- and hyperstomy.

Multiple lines of evidence indicate selection on stomatal ratio, but there is little consensus on the adaptive significance. Stomatal ratio varies widely, but nonrandomly [15, 17, 11, 19, 20, 21] and evolves rapidly in some taxa, possibly due to selection [22, 23, 24]. Several environmental and anatomical factors have been hypothesized to favour amphistomy (Table 1). The mechanistic details and literature underlying these hypotheses and predictions are described in Text S1. The preponderance of hypostomy almost certainly reflects a cost of upper stomata. For example, hypostomy has evolved anew in resupinate leaves [25]. Upper stomata might be costly because they increase susceptibility to foliar pathogens (e.g. rust fungi) that infect through stomata [13], suggesting that stomatal ratio mediates a tradeoff between photosynthetic rate and defence [23], but other costs have been proposed (Text S1). Identifying the selective forces (i.e. fitness benefits and costs) shaping stomatal ratio have been hampered by four methodological limitations. Namely, previous studies were typically qualitative rather than quantitative, confined to specific geographic regions or clades, did not account for phylogenetic nonindependence, and did not take into account multiple confounding factors. To overcome these limitations, I assembled a quantitative, global, and phylogenetically extensive database that disentangles correlated predictor variables (e.g. light level and leaf thickness).

**Table 1.**
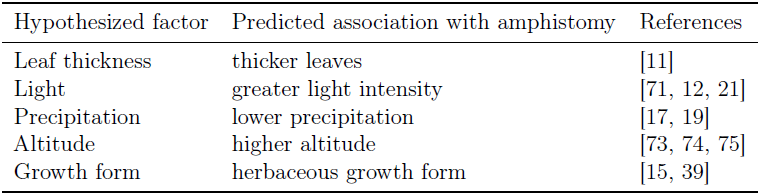
Adaptive hypotheses and predictions for stomatal ratio. The first and second columns indicate the hypothesized ecological factors and the predicted direction of association with amphistomy, respectively. References to key studies are provided, but see Text S1 for additional detail.

This new dataset revealed that stomatal ratio is a multimodal trait (Fig. 1). To test whether the observed pattern is consistent with constraint, I modified previous evolutionary process models to accommodate bounded traits like stomatal ratio. Fitting this model to the data indicates that stomatal ratio is highly constrained by a rugged adaptive landscape with multiple selective regimes (for discussion of selective regimes, see [26, 27, 5]). This led me to evaluate whether selection is sufficient to account for inferred constraints using theoretical and empirical approaches. First, I constructed a simple cost-benefit model consistent with the underlying physics and a minimum of additional assumptions. This model indicates that distinct peaks in the adaptive landscape can result from selective constraints, even when the underlying environmental gradients are smooth. In contrast, a review of the literature does not support a large role for genetic, developmental, and functional constraints. Finally, phylogenetic multiple regression identifies life history evolution as the primary selective agent underlying peak shifts, but anatomical and climatic factors are also important. By merging theory and data, this study adduces compelling new evidence that selection is the primary constraint on phenotypic evolution, at least for stomatal ratio. There is no reason to believe this trait is exceptional among functional traits, and hence the inferences drawn here could be generalizable to many other phenotypes that exhibit similar patterns indicative of evolutionary constraint.

**Fig. 1.**
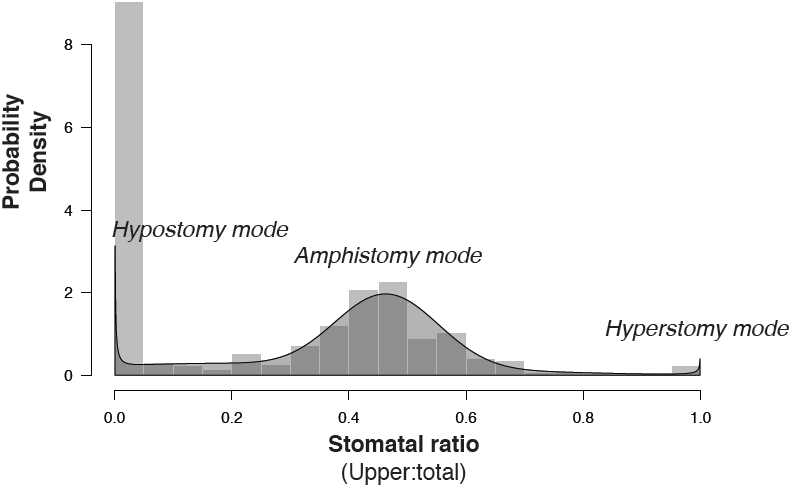
Stomatal ratio is a multimodal trait. A density histogram of stomatal ratio across 599 species (light grey bars in background) displays three noticeable modes. The plurality of species are completely hypostomatous (all stomata on the lower surface; stomatal ratio equals zero). There is a smaller, broader mode of amphistomatous species (approximately equal density of stomata on upper and lower surfaces; stomatal ration equals approximately one-half). Finally, there are a small number of hyperstomatous species (all stomata on the upper surface; stomatal ratio equals one). A mixture of selective regimes (shaded grey polygon) manifests these three modes, indicating that they are real features of constrained trait evolution rather than random noise.

## Results

### Stomatal ratio evolution is constrained by multiple selective regimes

I compiled a new, global dataset from 25 previously published studies (Text S2) containing trait data (stomatal ratio and leaf thickness) on 599 species across 94 plant families; the dataset with trait and climate data comprised a 552 species subset covering 90 families. The most striking feature of the data is that stomatal ratio (SR) is highly multimodal (Fig. 1), with apparent modes at 0 (hypostomatous), ≈ 0.5 (amphistomatous), and 1 (hyperstomatous). Note that here I am reporting stomatal ratio as the ratio of upper density to total density so that the distinct hypo- and hyperstomatous modes can be seen. Stomatal ratio does not conform to a nonmodal, uniform distribution (Komologrov-Smirnov test, *D* = 0.433, *P* = 1.11 × 10^−15^), even after removing all hypostomatous (SR = 0) species (K-S test, *D* = 0.293, *P* = 1.33 × 10^−15^). The data are also inconsistent with a unimodal, truncated exponential distribution bounded by 0 and 1 (K-S test, *D* = 0.429, *P* = 1.11×10^−15^).

In contrast, the distribution of stomatal ratio values across species is consistent with an evolutionary process model that includes constraints imposed by multiple selective regimes, indicating a rugged adaptive landscape. Although the results presented in this section only identify constraint, not necessarily selective constraint, I use selective regime because evidence in the following sections indicates that selection is the primary constraint. To infer regimes, I augmented a commonly used model of selective regimes, the Ornstein-Uhlenbeck process [28], to account for traits like SR that are bounded by 0 and 1 (see Materials and Methods and Text S3 for further detail and mathematical derivation). Under a bounded Ornstein-Uhlenbeck process model, the stationary distribution of stomatal ratio (or any proportion trait) *r* follows a Beta distribution:

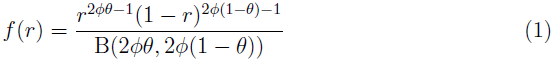

B(·) refers to the Beta function. A selective regime at stationarity is characterized by two parameters, a long-run average or ‘optimum’ in the adaptive landscape, *θ*, and a precision, *φ*, around the optimum. Greater values of *φ* produce distributions that are more tightly constrained around the optimum.

If a trait evolves on an adaptive landscape with multiple peaks, then a model with multiple selective regimes should fit the data better than a model with a single regime [27, 5]. I used finite mixture model analysis (Text S4) to estimate the number of selective regimes. This approach differs from conceptually similar methods, but can be applied to non-Gaussian traits like SR (see [29, 30] for alternative methods with Gaussian traits). I inferred three selective regimes (Table 2), but note that the mapping between modes and regimes is not always one-to-one. In particular, one regime produces modes at both 0 and 1 (Fig. S1). Nevertheless, the data clearly support the large number of hypostomatous (SR = 0) species as a distinct mode (Fig. S1). There was also strong support for an amphistomatous regime (compare Fig. S1A to Fig. S1B). Finally, the best-supported model also included a small mode for hyperstomatous species and a separate, smaller regime for species intermediate between hypo- and amphistomy (Fig. S1C).

**Table 2.** Multiple selective regimes are manifest in a multimodal trait distribution. Models with multiple components (*k*) corresponding to distinct selective regimes under a bounded Ornstein-Uhlenbeck process fit the data significantly better than models with a single regime (lower Bayesian Information Criterion [BIC]). In particular, the model with with three regimes is much more strongly supported than models with one or two regimes (see Fig. S1 for a visual representation of regimes). A mixture of multiple regimes, in turn, gives rise to a multimodal distribution with hypo-, amphi-, and hyperstomatous modes. For a given mixture, each of *k* regimes is represented as a component *i* parameterized by the strength of constraint ((*φ_i_*) around the long-term average (*θ_i_*) and a mixture weight *W_i_*.

| $k$ | Parameters | | | log-likelihood | df | BIC |
| --- | --- | --- | --- | --- | --- | --- |
| 1 | $\phi_1 = 0.4$ | $\theta_1 = 0.17$ | $w_1 = 1$ | -604 | 2 | 1220.9 |
| 2 | $\phi_1 = 0.25$ | $\theta_1 = 0.04$ | $w_1 = 0.52$ | -252.5 | 5 | 536.9 |
| | $\phi_2 = 9.98$ | $\theta_2 = 0.46$ | $w_2 = 0.48$ | | | |
| 3 | $\phi_1 = 0.16$ | $\theta_1 = 0.02$ | $w_1 = 0.47$ | -237.7 | 8 | 526.6 |
| | $\phi_2 = 17.24$ | $\theta_2 = 0.47$ | $w_2 = 0.38$ | | | |
| | $\phi_3 = 2.04$ | $\theta_3 = 0.35$ | $w_3 = 0.16$ | | | |
| 4 | $\phi_1 = 6.99$ | $\theta_1 = 0$ | $w_1 = 0.44$ | -235.6 | 11 | 541.6 |
| | $\phi_2 = 1.6$ | $\theta_2 = 0.35$ | $w_2 = 0.17$ | | | |
| | $\phi_3 = 16.85$ | $\theta_3 = 0.47$ | $w_3 = 0.38$ | | | |
| | $\phi_4 = 181.8$ | $\theta_4 = 0.99$ | $w_4 = 0$ | | | |

The same general pattern seen at the global scale – multiple selective regimes leading to distinct modes – is recapitulated within nine of ten families best-represented in the global dataset (Fig. 2). Two regimes are supported in most (8 of 9) multi-regime families, except Asteraceae, in which three regimes are favoured (Fig. 2A). In one family, Rubiaceae, all species were inferred as members of a hypostomatous regime. In all mutli-regime families except Poaceae, there are distinct regimes associated with hypo- and amphistomy; in Poaceae, there are hyper- and amphistomous regimes instead (Fig. 2E). However, the hyperstomatous species of Poaceae in this study may not be representative of family since they are wetland specialists in the genus *Spartina* [31]. Generally, the internal (i.e. amphistomatous) mode is closely centered around 0.5, as predicted from biophysical theory [11, 13], except in in the Rosaceae, where the inferred optimum is closer to 0.25. Although I was unable to account for phylogenetic nonindependence in these analyses (see Materials and Methods), that a similar pattern – species are either amphistomatous or hypo/hyperstomatous, but rarely intermediate – emerges independently in multiple families indicates the conclusions are unlikely to change qualitatively once fully phylogenetic methods can be extended to bounded traits. In summary, the apparent pattern of constraint on stomatal ratio is strikingly similar across multiple disparate families and at a global scale, suggesting convergent evolution because of shared phenotypic constraint.

**Fig. 2.**
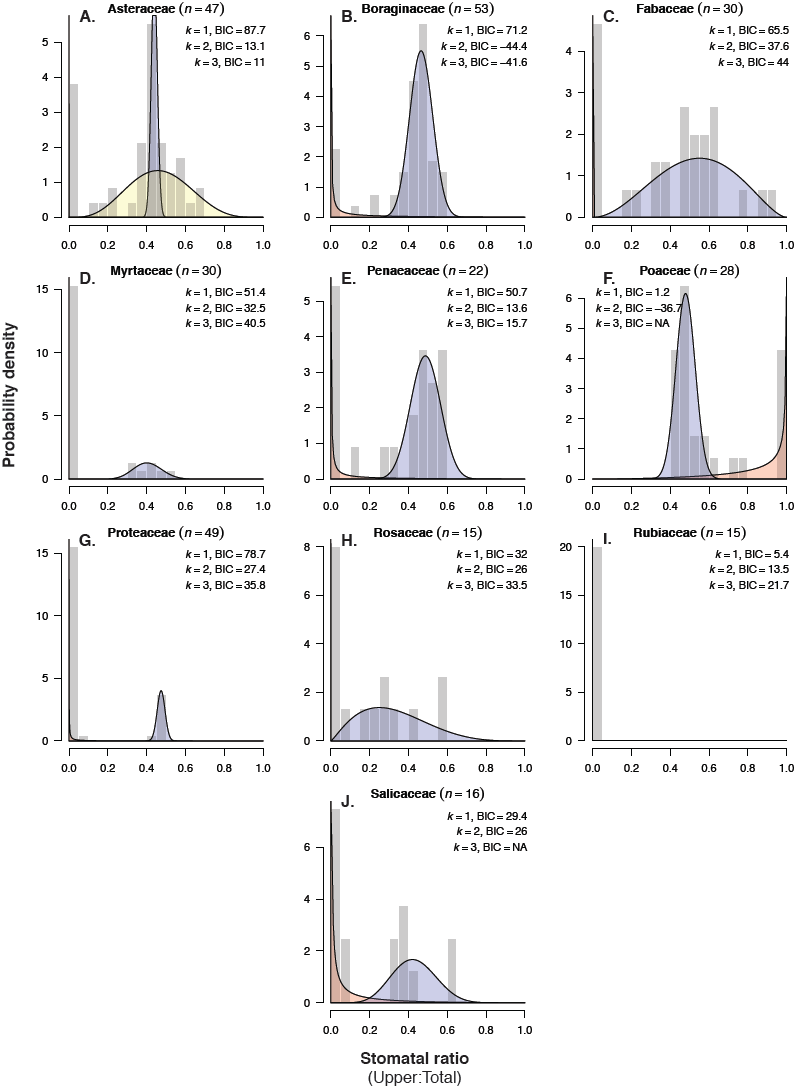
Repeated evolution of multimodality suggests that the adaptive landscape is conserved across land plants. Shaded polygons of inferred regimes are plotted atop a histogram (grey bar) of stomatal ratio from a given plant family (grey bars). Note that some distributions are very narrow spikes near the origin. The title gives the family name and number of species sampled *n* from that family. Three regimes were inferred for Asteraceae (panel **A**.); two regimes were inferred for other families except the Rubiaceae (panels **B.-J.**). The number of regimes was inferred from information theoretic comparisons of finite mixture models with Beta-distributed components. The Bayesian Information Criterion (BIC) for models with *k* = 1, 2, and 3 components is given in the top. I accepted models with additional regimes (higher *k*) if they decreased BIC by two or more. In Poaceae and Salicaeae, I rejected models with *k* = 3 because some components had very low membership.

### Selection is sufficient to accommodate constraint

I analyzed a simple cost-benefit model of stomatal ratio to ask whether selection is sufficient to account for apparent phenotypic constraint. Not surprisingly, selection favours greater stomatal ratio (*S*_fit_) as the fitness benefit of greater photosynthesis increases relative to the cost of upper stomata (Fig. 3A-C), but the shape of the function is highly sensitive to one parameter in the model, *σ*^2^. In particular, the adaptive landscape goes from being smooth when *σ*^2^ is high to rugged when *σ*^2^ is low (Fig. 3D-F). When the landscape is smooth, intermediate phenotypes between complete hypostomy and amphistomy are best when the benefit:cost ratio itself is intermediate. In contrast, when the landscape is rugged, intermediates are universally less fit than either of the boundary phenotypes. In a rugged landscape, as the benefit:cost ratio decreases there is a sudden shift from amphistomy being favoured to hypostomy being favoured. The dearth of species with intermediate *SR* in nature, especially within families, therefore suggests that the adaptive landscape for stomatal ratio is generally rugged. Numerical simulations based on smooth variation in the benefit:cost ratio indicate that the simple, yet realistic assumptions of this model are sufficient to generate qualitatively similar patterns of multimodality to those seen in nature (Fig. 3G-H).

**Fig. 3.**
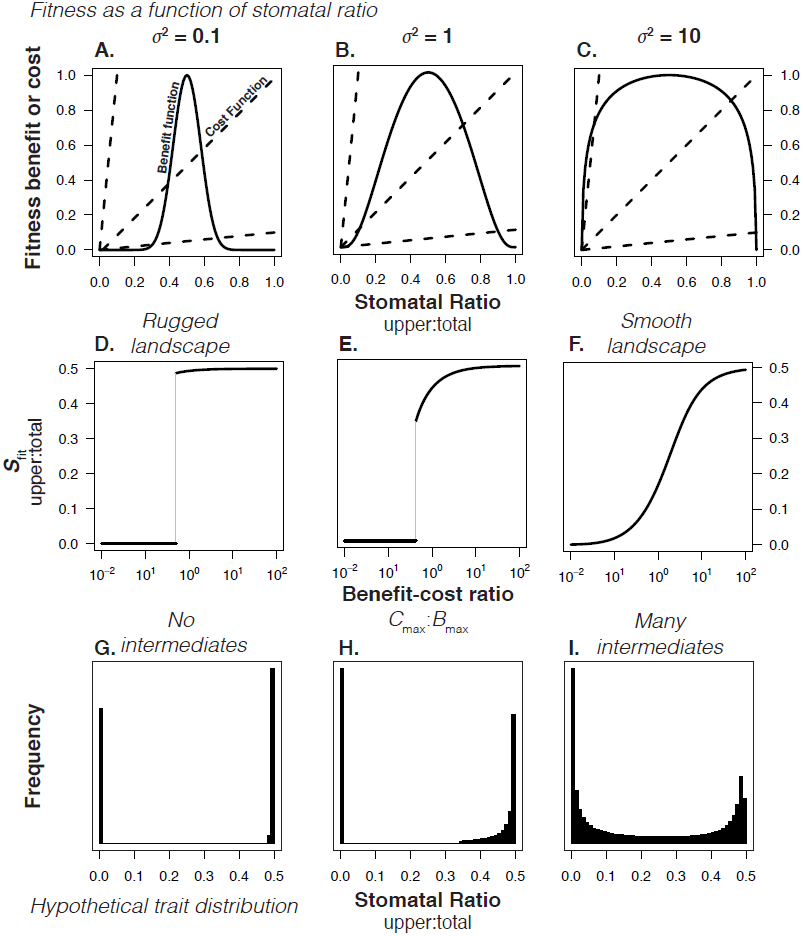
Selection is sufficient to explain why intermediate phenotypes are universally unfit and the adaptive landscape is rugged. Panels **A.-C:** In each panel, a benefit function (solid line, see Eq 3) is shown with three different cost functions (dashed line, see Eq 5). In all panels, *B*_max_ is fixed at 1 and three slopes of the cost function, *C*_max_ are illustrated: 0.1 (shallow slope), 1 (medium slope), and 10 (steep slope). The fitness benefit is always maximized when stomatal ratio is 0.5 (amphistomy), corresponding to *S*_opt_ = 0 on a logit scale. The shape factor *σ*^2^ changes the benefit function from bell-shaped in **A**. to an inverted-U shape in **C**. Panels **D.-F**. show that the shape of the benefit function affects the topography of the adaptive landscape. Solid lines are the stomatal ratio that optimizes fitness (*S*_fit_) as a function of the benefit:cost ratio (*B*_max_ : *C*_max_). When the benefits are high compared to costs, amphistomy (stomatal ratio = 0.5) is favoured; when the costs are high, hypostomy is favoured (stomatal ratio = 0). However, the transition between these extremes can be abrupt when the landscape is rugged (panel **D**.) or gradual when the landscape is smooth (panel **F**.). The light gray line indicates the range of universally unfit phenotypes. Panels **G.-I.** show hypothetical trait distributions assuming that the benefit:cost ratio varies uniformly from 10^−2^ to 10^2^. Histograms were generated by solving for *S*_fit_ with 10^4^ evenly spaced values of *B*_max_ : *C*_max_. Note that the trait values range from hypostomatous to amphistomatous (stomatal ratio = 0.5), but a mirror image distribution with hyperstomatous species would be seen if fitness costs accrued to lower stomata.

### Growth form, leaf thickness, and precipitation shape stomatal ratio evolution

If stomatal ratio is strongly associated with other traits or climatic factors, especially if there are compelling *a priori* hypotheses (Table 1) supporting such associations, then it suggests that trait variation is shaped by selection. Phylogenetic multiple regression consistently identified growth form and, to a lesser extent, leaf thickness and precipitation as the best predictors of stomatal ratio (Table 3). Amphistomy was strongly associated with fast growth forms (herbaceous plants), whereas hypos-tomy was most common in slower growing shrubs and trees (Fig. 4). As predicted by biophysical theory [11, 13], thicker leaves also tended to be amphistomatous, although the correlation was weak (Fig. S2A). Finally, amphistomy was more common in dry environments, whereas hypo/hyperstomy were associated with higher precipitation (Fig. S2B). Elevation and leaf area index, a proxy for open habitat, were not significantly associated with stomatal ratio in this dataset (Table 3). In single regressions, amphistomy was more common more open environments, as in previous studies [12, 18, 19, 21], but this correlation was not significant after precipitation was factored into multiple regression (precipitation and leaf area index are positively correlated).

**Table 3.** Growth form, anatomy, and precipitation jointly determine stom-atal ratio. Three models with varying levels of phylogenetic signal (Brownian motion [top], Pagel’s λ [middle], and nonphylogenetic [bottom]) identify growth form, leaf thickness, and mean annual precipitation as significantly associated with stom-atal ratio.

| Stomatal Ratio ∼ | df | SS | MS | F | $P$ |
| --- | --- | --- | --- | --- | --- |
| *Brownian Motion* |  |  |  |  |  |
| log(Leaf Thickness) | 1 | 0.017 | 0.017 | 20.31 | $8.08 \times 10^{-6}$ |
| Mean Annual Precipitation | 1 | 0.021 | 0.021 | 24.11 | $1.21 \times 10^{-6}$ |
| Elevation | 1 | 0 | 0 | 0.08 | 0.78 |
| Leaf Area Index | 1 | 0 | 0 | 0.05 | 0.82 |
| Growth Form | 5 | 0.039 | 0.008 | 9.06 | $2.74 \times 10^{-8}$ |
| *Pagel’s $\lambda = 0.64$* |  |  |  |  |  |
| log(Leaf Thickness) | 1 | 0.008 | 0.008 | 24.38 | $1.05 \times 10^{-6}$ |
| Mean Annual Precipitation | 1 | 0.009 | 0.009 | 26.03 | $4.67 \times 10^{-7}$ |
| Elevation | 1 | 0 | 0 | 0.26 | 0.61 |
| Leaf Area Index | 1 | 0 | 0 | 0 | 1 |
| Growth Form | 5 | 0.027 | 0.005 | 15.52 | $2.77 \times 10^{-14}$ |
| *Nonphylogenetic* |  |  |  |  |  |
| log(Leaf Thickness) | 1 | 2.376 | 2.376 | 31.67 | $2.94 \times 10^{-8}$ |
| Mean Annual Precipitation | 1 | 1.711 | 1.711 | 22.81 | $2.31 \times 10^{-6}$ |
| Elevation | 1 | 0.009 | 0.009 | 0.12 | 0.72 |
| Leaf Area Index | 1 | 0.031 | 0.031 | 0.41 | 0.52 |
| Growth Form | 5 | 15.897 | 3.179 | 42.38 | $7.36 \times 10^{-37}$ |

**Fig. 4.**
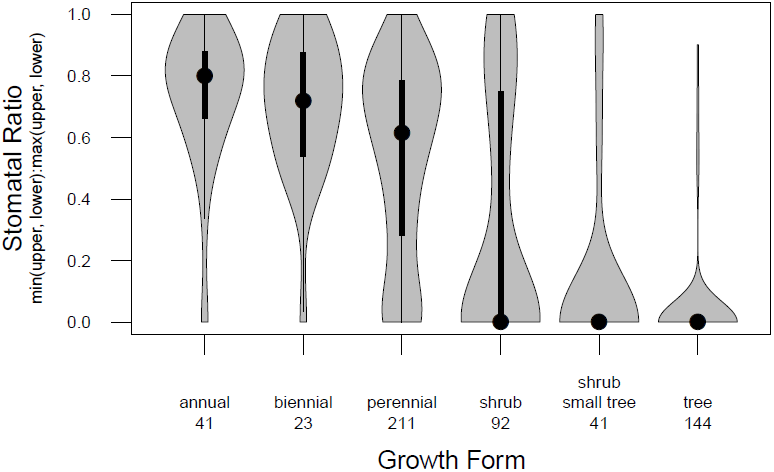
Association between amphistomy and fast growth forms points to selection on life history shaping stomatal ratio evolution. The violin plot shows of stomatal ratio as a function of growth form across all species in the dataset. The width of the grey polygons indicates the density of data. Length of grey polygon indicate the range of the data; the point indicates the median; the thick lines indicate the 0.25 and 0.75 quantiles. Sample sizes per growth form in the dataset are given below the label.

## Discussion

Phenotypic evolution is often constrained, but the relative role of selective versus nonselective constraints is unclear. This study posits that multimodal traits reveal distinct peaks of high fitness in a rugged adaptive landscape. Hence, the prevalence of certain phenotypes and the dearth of others directly reflects selective constraints on phenotypic evolution. Evidence from a new, global dataset clearly shows that stomatal ratio is a multimodal trait (Fig. 1) and that multimodality has evolved repeatedly in land plants (Fig. 2). These patterns are difficult to reconcile with models omitting constraint, but are consistent with a rugged adaptive landscape comprised of multiple selective regimes (Table 2). A simple cost-benefit model of stomatal ratio shows that selection is a sufficient explanation, particularly when the underlying adaptive landscape is predominantly rugged. Adaptive evolution from one peak in the landscape to another (i.e. hypo- to amphistomy or *vice versa*) appears to be primarily driven by growth form, suggesting that the fitness benefit of amphistomy – faster diffusion of CO_2_ to chloroplasts – is greatest in species with ‘fast’ life histories.

### Multimodality implies constraint on the macroevolutionary adaptive landscape

Just as water is only found as ice, liquid, and steam, despite continuous variation in temperature, stomatal ratio comes in partially discrete clusters corresponding to hypo-, amphi-, and hyperstomy, but less often intermediate (Fig. 1). In fact, the modes identified here correspond remarkably with traditional botanical classifications [32], suggesting that these workers recognized the pattern even without quantitative analyses. The multimodal pattern in the dataset cannot be explained by an evolutionary process model neglecting constraint (Text S3). However, apparent clustering could occur by systematic underrepresentation of intermediate trait values [33] or nonrandom taxon sampling. It is highly improbable that intermediate phenotypes exist at greater frequency in nature but are rarely reported, as most studies have no *a priori* hypothesis about stomatal ratio in their study organisms. If anything, by omitting many studies that report only qualitative data, I might have enriched the frequency of intermediate phenotypes, as these are the most likely to be reported quantitatively. Nonrandom taxon sampling, without accounting for phylogeny, could also give the appearance of multimodality. To give an extreme example, if there had been a single transition from hypo- to amphistomy followed by stasis, then sampling the tips of the phylogeny would produce a multimodal pattern with apparently strong statistical support, even though it only represents a single evolutionary event. Methodological limitations prevented me from fully accounting for phylogeny (see Materials and Methods), but the fact that multimodality reappears in multiple distantly-related families (Fig. 2) makes nonrandom taxon sampling alone an unlikely explanation, though it might accentuate the pattern. Future work is needed to extend regime-inference methods [27, 29, 30] to non-Gaussian traits, as this study begins to do with a new evolutionary process model for proportion traits.

### Selection is the most likely explanation for phenotypic constraint

In principle, constraint could reflect a mix of selective, genetic, developmental, and functional factors [7]. However, the preponderance of available theory and data on stomatal ratio suggests selection is responsible for most if not all of the phenotypic constraint. Genetic, developmental, and functional constraints cannot explain the dearth of intermediate phenotypes because intermediates are genetically accessible as well as developmentally and functionally possible. The appropriate mutations to generate intermediate phenotypes occur spontaneously during mutagenesis [34], segregate among natural populations [35, 36, 37, 23], and are fixed between closely related species [38, 24].

In contrast, the cost-benefit model presented here shows that with a small number of realistic, evidence-based assumptions, selection is sufficient to accommodate the data and helps clarify why discrete modes form even when the underlying environmental gradients are smooth (environmental gradients need not be smooth, but it is unnecessary to assume otherwise). Stomata are often distributed equally on both surfaces (amphistomy) because this arrangement optimizes photosynthetic rate. This was an assumption of the model based on biophysical theory [11, 13]. More often, all stomata are on the lower surface because the costs of upper stomata outweigh the benefits. A dearth of intermediates between hypo- and amphistomy occurs when the landscape is rugged, making these phenotypes often fall in a fitness valley. However, the best mixture model includes a small peak of these intermediates (Table 2, Fig. S1). This suggests that although the adaptive landscape is constrained and often rugged, it may shift from rugged to smooth over macroevolutionary time. However, the fact that most species, especially within families (Fig. 2), cluster around particular modes suggests that the landscape is predominantly rugged. Finally, the small number of hyperstomatous species indicates that there are occasionally situations in which upper stomata are favoured, such as in aquatic plants or those with unusual epidermal or spongy mesophyll anatomy.

### Life history, more than anatomy and climate, determines stom-atal ratio

Nonrandom association between stomatal ratio, other ecologically important traits, and climate also supports a significant role for selection in shaping trait evolution. To my knowledge, this is the first study to rigorously demonstrate a strong association between growth form and stomatal ratio, although it had been suggested by earlier ecological surveys [15, 39]. Two hypotheses that might explain the relationship between growth form and stomatal ratio are: 1) herbaceous plants have shorter leaf lifespans [40], requiring higher photosynthetic rates to pay their construction costs in a shorter time [41]; 2) herbaceous plants have faster life histories, leading to stronger selection on high growth rates, mediated in part by higher leaf-level photosynthetic rate [42]. That the relationship between stomatal ratio and whole-plant lifespan holds within herbaceous (annuals vs. perennials) and woody (shrubs vs. trees), supports the second hypothesis (selection on faster life history favours amphistomy). Although this hypothesis requires further testing, if correct, it implies remarkably strong selection on leaf-level photosynthesis, as the photosynthetic advantage of amphistomy over hypostomy is only a few percent in a typical herbaceous leaf [11].

Surprisingly, I found little evidence supporting the most common adaptive explanation for amphistomy, that thicker leaves ‘need’ stomata on both sides to facilitate CO_2_ diffusion [11]. In actuality, support for this hypothesis is mixed (Text S1), especially when phylogenetic nonindependence is taken into account [43, 39] (but see [44]). It is now clear why previous studies came to different conclusions: thicker leaves do tend to be amphistomatous, even once phylogeny is accounted for, but the trend is weak (Fig. S2A). Less powerful studies than this one could easily have failed to detect a significant relationship. Hence, leaf thickness, by constraining CO_2_ diffusion, imposes selection for amphistomy. I also found that amphistomy was more common in plants from low precipitation environments. For a given stomatal conductance, which is proportional to evaporative water loss, amphistomy improves water-use efficiency by increasing photosynthetic rate [11], suggesting a plausible mechanism for selection on amphistomy in dry environments. Although low precipitation was correlated with habitat openness, measured using leaf area index, multiple phylogenetic regression indicated that precipitation was causal, in contrast to previous studies [18, 21]. These studies used finer scale (plant-level) descriptions of light environment that might have been missed by the coarser, satellite-based measurements of canopy cover used here. Alternatively, patterns at the global scale might differ from those within particular families or biomes. Finally, I was unable to test the effects of leaf orientation and stomatal packing on stomatal ratio, though these are likely to be important factors in many plants [20]. The evidence from this and previous studies shows that stomatal ratio is an ecologically relevant functional trait that could be valuable in physiological ecological and evolution [45].

That many ecologically important traits, like stomatal ratio, cluster around particular values but not others suggests pervasive constraint on phenotypic evolution. How can we seek a general explanation for this pattern when any particular instance requires specific mechanistic and ecological knowledge about a focal trait? For example, the emerging evidence from this and other recent studies on stomatal ratio (see especially [23]) is that peaks of high fitness are constrained by a tradeoff between photosynthetic rate and defence against foliar pathogens that preferentially infect though upper stomata. In particular, the cost-benefit model analyzed here predicts that even a small change in the fitness costs or benefits are sufficient to shift fitness peaks into qualitatively different selective regimes. If it is generally true that multimodal traits are associated with rapid regime shifts, then one way forward is to look for signatures of such shifts in closely-related species that sit astride different regimes. For example, one signature of regime shifts could be the presence of quantitative trait loci large enough to pass over valleys separating fitness peaks. Consistent with this, [24] recently identified two large effect loci that together are capable of making a hypostomatous leaf amphistomatous, perhaps suggesting that these loci enabled a regime shift. Integrating comparative biology, mechanistic studies of organismal function, and the genetics of adaptation, as this and others studies [46] have begun to do, points to a general approach for evaluating the common features of macroevolutionary adaptive landscapes and, hence, the role of selection in constraining phenotypic evolution.

## Materials and Methods

### Assembling a comparative data set

#### Stomatal ratio and leaf thickness

I collected quantitative data on stomatal ratio and leaf thickness from previously published studies (see Text S2 for full list of sources). These data are spread across a large and diverse literature, including functional ecology, taxonomy, agriculture, and physiology. Hence, neither a standardized nor exhaustive search was possible. I started by using Web of Knowledge to locate studies that cited seminal papers on the adaptive significance of amphistomy, specifically [11] and [12]. Once I found a paper with data, I examined papers that cited those ones. Finally, I found additional data sources in comprehensive reviews of plant anatomy [47, 32, 48]. For all data papers, I recorded the mean leaf thickness, abaxial (lower) and adaxial (upper) stomatal density for each species. Where only ranges were given, I used the midpoint. If the study included a treatment, I collected only data from the control treatment. If studies measured both juvenile and adult leaves, I used only adult leaves (no study reported only juvenile leaves). Usually data were given in a table, but occasionally I used ImageJ [49] to extract data from figures or contacted authors for data. I only included data from studies that intentionally examined both surfaces for stomata; I excluded data from studies that described species categorically as “hypostomatous”, or “amphistomatous”, or “hyper-stomatous”. Excluding qualitative data was necessary because there is no standard definition of “amphistomy” – it has sometimes been used to describe species that have approximately equal densities on each side [11] and at other times for species that have any stomata on the both surfaces [16, 15].

#### Climate and elevation

Based on the *a priori* hypotheses, I extracted data on mean annual precipitation (average 1950 – 2000), elevation (Worldclim [50]), and light environment (average leaf area index between 1982 – 1998 based on remote sensing [51]). For light environment, I used a satellite indicator of leaf area index, the number of leaf layers between the ground and top of the canopy. Lower leaf area index is interpreted as a more open light environment. The strength of these global data sources is that I was able to obtain data for every species from the same dataset. A limitation of these data is that even the highest resolution (≈ 1 km) data might miss important temporal and microsite variation. I discuss these limitations in light of the findings in the Discussion. For climate and elevation, geographic coordinates for each species are needed. For this, I downloaded all georeferenced herbarium specimens for a given species from GBIF (last accessed Jan 15–18, 2015) using the occ_search function in rgbif [52]. I filtered out or manually edited clearly erroneous locations (e.g. lat = 0 or lon = 0 or where lat and lon were clearly reversed). The mean and median number of GBIF georeferenced occurrences per species was 737 and 194, respectively. I calculated the trimmed-mean (10% trim) mean annual precipitation, elevation, and leaf area index to further remove specimens well outside the species’ range, possibly because they were, say, misidentified, cultivated, or improperly georeferenced.

#### Growth Form

I partitioned species by growth form into the following categories: trees, small trees/shrubs, shrubs, and herbaceous species (forbs and grasses). Herbaceous species were further subdivided into annuals, biennials, and perennials. Species that were variable or intermediate (e.g. annual/biennial, annual/perennial, biennial/perennial, or annual/biennial/perennial) were classified as ‘biennial’. Subshrubs with some woody growth were lumped with perennials rather than shrubs. Where possible, I obtained growth form data from associated data papers. When this information was not given, I used regional floras, supplemented by online trait databases such as USDA Plants [53] and Encyclopedia of Life [54]. When these sources were unavailable or ambiguous for a given species, I checked the primary taxonomic literature by searching the species name in Google Scholar.

##### Taxonomic name resolution

I submitted taxonomic names in the database to the Taxonomic Name Resolution Service (TNRS) [55]. I used names given by TNRS when it returned an accepted name or synonym with overall score greater than 0.97 (scores are between 0 to 1). I scrutinized names where TNRS deemed the name illegitimate, gave no opinion, or was otherwise ambiguous. At that point, I consulted additional plant taxonomic repositories: The Plant List [56], International Plant Names Index [57], and the Euro+Med PlantBase [58]. When no accepted names were identified, I used original name given by the authors. For two very recent papers [59, 60], I used the names given by those authors.

### Pattern to process: connecting multimodality to phenotypic constraint

Comparative methods often infer constraint by comparing the fit of evolutionary process models with and without constraint. Constraint, usually interpreted as a selective regime, is typically modelled as an Ornstein-Uhlenbeck process [28, 27, 5], but this model is inappropriate for proportion traits like stomatal ratio. I therefore developed a new evolutionary process model that is analogous to an Ornstein-Uhlenbeck process except that traits are bounded by 0 and 1. A full description of model assumptions and a derivation of the stationary distribution under a given selective regime are available in Text S3. The key result is that a trait evolving under a single selective regime should conform to a Beta distribution at stationarity.

Multimodality suggests the presence of multiple selective regimes associated with different modes. I tested for multiple regimes using a conceptually similar but somewhat different approach than previous studies. Current methods for inferring multiple selective regimes are in their infancy [27, 29, 30] and cannot yet accommodate Beta-distributed traits because I could not obtain a general solution to the stochastic differential equation in Text S3. Future work is needed to develop numerical methods, such as Approximate Bayesian Computation [61], to integrate the bounded Ornstein-Uhlenbeck process model elaborated here into existing statistical frameworks for multi-regime inference. However, a few lines of reasoning I discuss below indicate that the main conclusions of this study are robust.

I used finite mixture models to infer the number of selective regimes shaping stomatal ratio evolution (see [6] for a similar approach). That is, I assume the current distribution of trait values across species can be represented as a mixture of multiple selective regimes at stationarity, each of which is modelled as a Beta-distributed variable. To fit models, I used an expectation-maximization algorithm to find the maximum likelihood mixture model from the data. A complete derivation of the likelihood function and a description of the fitting algorithm are given in Text S4. R code to implement the algorithm is available on Dryad [62]. I selected the best model using the more conservative Bayesian Information Criterion (BIC) to compensate for the fact that I am not accounting for phylogenetic nonindependence in this analysis (see below). I accepted models with an additional selective regime if they decreased BIC by 2 or more. By fitting the data to the stationary distribution, I implicitly assume that evolution is sufficiently rapid to ignore phylogenetic signal. Numerical simulations of the diffusion indicate that the transitory distribution is also Beta (data not shown), meaning that evidence for multiple regimes (i.e. a better fit of a mixture model with multiple Beta components) cannot be an artifact of transitory behaviour within a single regime. I also tested for multiple regimes within families where there was sufficient data (*n* ≥ 15). Ten families met this criterion. For each family, I compared the fit of mixtures with *k* = 1, 2, or 3 regimes, accepting models with an additional regime if they decreased BIC by 2 or more. Further, I rejected additional regimes supported by BIC if one of those regimes contained fewer than 3 species (this affected Poaceae and Salicaceae). Although testing for multiple regimes within families using the stationary distribution is an imperfect substitute for fitting the process model to the entire tree, it is nevertheless informative. If multiple regimes are found repeatedly in disparate families, this provides compelling evidence for convergent evolution because of phenotypic constraints imposed by similar adaptive landscapes.

### Is selection sufficient to account for multimodality?

In this section, I use theory to ask under what conditions selection can explain the rugged adaptive landscape implied by fitting the evolutionary process model to the data. First, I ask whether a model with simple fitness costs and benefits of upper stomata produces multiple fitness peaks (Text S1 discusses the fitness benefits and costs associated with stomatal ratio). Next, I examine whether such a landscape generates a trait distribution that qualitatively resembles the data, even when the underlying environmental gradients are smooth. I specifically focus on the pattern observed within families, where there was generally one mode of amphistomatous species and another mode of hypostomatous species (hyperstomatous in the case of Poaceae). I also opted to tradeoff the precision of a biophysical diffusion model for a more general, albeit realistic, model with fewer parameters. Hence, the cost-benefit model of stomatal ratio is true to the underlying physics but otherwise not strongly dependent on specific assumptions. Future work will be needed to test if this more general model is consistent with mechanistic biophysical models. The symbols used in the model are summarized in Table 4.

**Table 4.** Glossary of symbols used in the cost-benefit model.

| Symbol | Description |
| --- | --- |
| $SR$ | Stomatal ratio: ratio of upper to total stomatal density |
| $S$ | logit of stomatal ratio ($SR$) |
| $S_{\text{opt}}$ | Stomatal ratio (logit scale) that maximizes fitness benefits |
| $B_{\text{max}}$ | Maximum fitness benefit when $S = S_{\text{opt}}$ |
| $\sigma^2$ | Shape factor of benefit function |
| $C_{\text{max}}$ | Maximum fitness cost of when all stomata are on the upper side ($SR = 1$) |
| $S_{\text{fit}}$ | Stomatal ratio maximizes fitness (benefits minus costs) |

I model selection on the logit of stomatal ratio (upper:total), which I denote *S* = logit(*SR*) = log (*SR*/(1 − *SR*)), so that feasible trait variation (*SR* is constrained from 0 to 1) is continuous and unbounded. Fitness as a function of stomatal ratio depends on the difference between the benefits (*f*(*S*)) minus the costs (*g*(*S*)). Therefore, fitness as a function of stomatal ratio is:

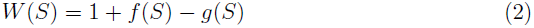

Based on biophysical theory [11, 13], I assume that there is an intermediate optimal stomatal ratio (*S_opt_*) at which photosynthetic rate is maximized. Above and below that optimum, photosynthetic rate decreases, which I modelled as a Gaussian function:

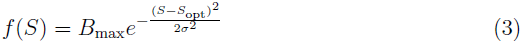

*B*_max_ defines the maximum fitness when *S = S_opt_*. *σ*^2^ acts akin to a shape factor when the function is viewed from a logit scale. When *σ*^2^ is large, the benefit function has an inverted-U shape. There are increasing returns to fitness of the first few upper stomata, but diminishing returns to further increases in SR (Fig. 3A). In contrast, when *σ*^2^ is small, the benefit function is more bell-shaped; the fitness benefit of the first few upper stomata is large, but with diminishing returns (Fig. 3C).

I assumed a linear cost (e.g. increased susceptibility to foliar pathogens [23]) for each additional upper stomate. The total cost as a function of stomatal ratio is the product of the total stomatal density, the stomatal ratio (upper:total density), and the cost per upper stomate. I define the slope of the cost function as *C*_max_, which is equal to the total stomatal density times the cost per upper stomate:

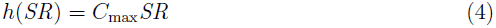

On a logit scale, the total cost asymptotically approaches *C*_max_:

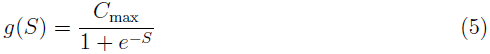

If more were known about the cost of having upper stomata, a more realistic model could be constructed. Without such knowledge, I believe it is judicious to start with the simplest model that makes few assumptions and therefore could apply to a large number of particular underlying mechanisms. Substituting Eqs 3 and 5 into Eq 2, fitness as a function of *S* is:

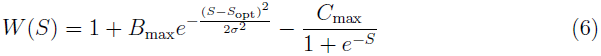

Note that if the cost function were applied to lower rather than upper stomata, as might be the case for specialized taxa such as aquatic plants, then one could obtain the same results, except that hyper- rather than hypostomy would prevail, as in the Poaceae data. The fitness function is maximized where the marginal benefit of the next upper stomate is equal to the marginal cost:

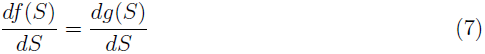

I did not obtain an analytical solution, so instead I used the optim function in R [63] to numerically solve for the stomatal ratio that maximized fitness (*S*_fit_) under varying ratios of fitness cost (*C*_max_) to benefit (*B*_max_). I tuned the benefit:cost ratio by fixing *B*_max_ to 1 and varying C_max_ between 0.01 and 100. I also varied the shape factor *σ*^2^ between 0.1 and 10, which appeared to capture the full range of relevant model behaviour. For all numerical solutions, I assumed that the optimal stomatal ratio for photosynthesis was 0.5, hence *S*_opt_ = 0 on a logit scale. Next, I generated hypothetical trait distributions under a scenario where the benefit:cost ratio varies uniformly from 10^−2^ to 10^2^. I solved for *S*_fit_ with 10^4^ evenly spaced values of *B*_max_ : C_max_ under low, medium, and high values of *σ*^2^. R code for finding numerical solutions is available from Dryad [62].

### Testing adaptive hypotheses for stomatal ratio using phylogenetic regression

I tested for an association between stomatal ratio, leaf thickness, mean annual precipitation, elevation, leaf area index, and growth form using type 2 phylogenetic ANOVA with both categorical (Growth form) and continuous (e.g. leaf thickness) predictor variables. For this analysis I quantified stomatal ratio as min(upper density, lower density):max(upper density, lower density). In this form, stomatal ratio equals 1 when the densities on each surface are the same, and goes to 0 as the distribution become more asymmetrical (hypostomy or hyperstomy). Note that this form differs from what I use in analyzing multimodality because I wanted to specifically test which factors favour the phososynthetically optimal distribution (amphistomy) versus suboptimal distributions (either hypo- or hyperstomy). I accounted for phylogeny using a Phylomatic [64] megatree for this relatively large and phylogenetically extensive dataset. To examine whether results were robust to phylogenetic correction, I analyzed the data using three methods: Brownian motion (high phylogenetic signal), Pagel’s λ (intermediate phylogenetic signal), and no phylogenetic signal (normal ANOVA). For the intermediate signal model, I estimated Pagel’s λ using maximum likelihood. Phylogenetic models were fit using phylogenetic least squares in the R package ‘caper’ [65].The trait dataset and phylogeny used in these analyses are available on Dryad [62].

## Acknowledgements

Ranessa Cooper, Jenny Read, Gregory Jordan, and Tim Brodribb generously made data available. Members of the Angert and Schluter labs provided feedback on an earlier version of this manuscript. I am supported by a Biodiversity Postdoctoral Fellowship funded by NSERC-CREATE.

## Supporting Information

**Fig. S1.**
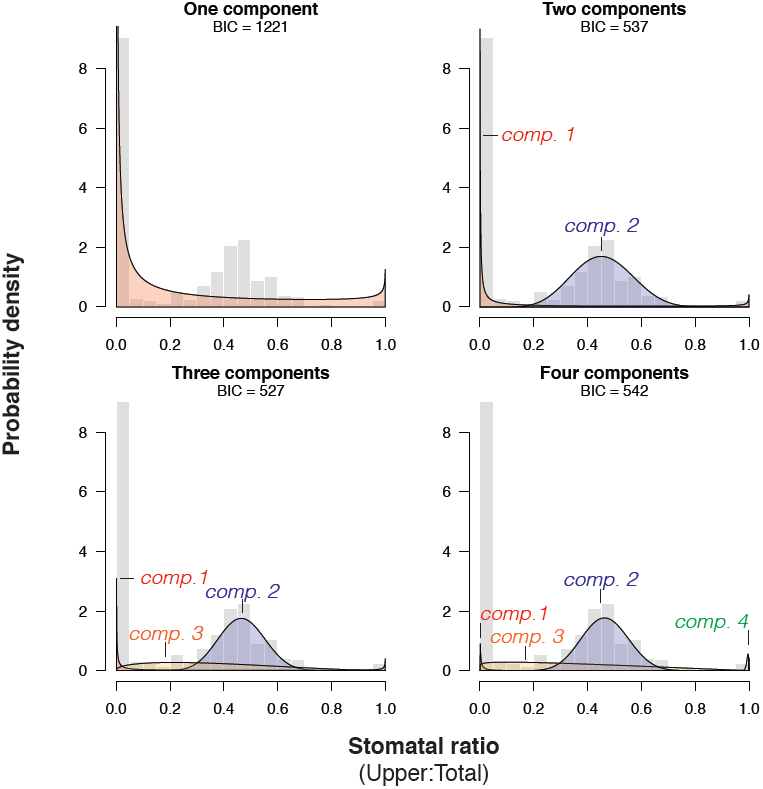
A multimodal trait distribution implies multiple selective regimes. Model selection using Bayesian Information Criterion (BIC) favoured models that were mixtures of composed multiple selective regimes. **A**. A model with one component was a poor fit because it cannot account for the large peak of amphistomatous species. **B**. A model with two components fit the data much better because it incorporates separate selective regimes for amphistomatous species (blue polygon) and hypo-/hyperstomatous species (red polygon). **C**. An additional selective regime (orange polygon) for species with stomatal ratios between 0 and 0.5 improved model fit, suggesting that intermediate phenotypes are favoured in some circumstances. **D**. Finally, a model with a fourth component (green polygon) did not significantly improve the fit (higher BIC).

**Fig. S2.**
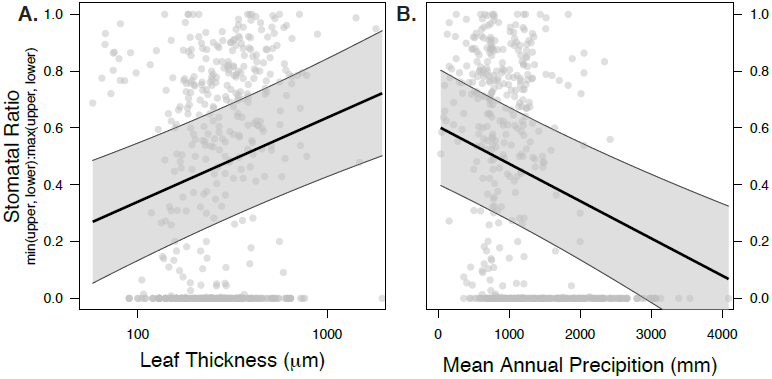
Amphistomy is weakly associated with thicker leaves and drier habitats. Each point represents a species from the global dataset. The thick line and gray polygon are the median and 95% confidence intervals from the posterior distribution of predicted stomatal ratio as a function of leaf thickness based on phylogenetic regression. The fitted lines and confidence intervals are drawn with growth form set to perennial and other continuous predictor variables set to their median.

### Text S1: Hypothesized benefits and costs of amphistomy

There are at least seven viable, non-mutually exclusive hypotheses for on the adaptive significance of amphistomy, five of which I evaluate here.

#### H1: Leaf thickness

The most widely cited and frequently tested diffusional limitation hypothesis is that amphistomy is adaptive in thick leaves. Models [11, 13] and experiments [14] demonstrate that the path length from substomatal cavities to chloroplasts can impose a large constraint on photosynthesis, especially when leaf thickness exceeds approximately 300 *µ*m. Several studies have found a positive correlation between leaf thickness and amphistomy [11, 20, 66, 67, 68, 59, 44], but the evidence is equivocal [69, 12, 43].

#### H2: Light

A second hypothesis is that amphistomy is favoured in high light, open environments because CO_2_ becomes more limiting at high irradiance. H1 and H2 are difficult to disentangle, and could even reinforce one another, because leaf thickness increases under high irradiance [70]. However, several studies have argued that the light environment, rather than leaf thickness, is the primary factor affecting selection on amphistomy [19, 71, 18, 12, 20, 21].

#### H3: Precipitation

Wood [17] observed that amphistomy was common in Australian deserts. Although amphistomy is sometimes common in dry environments, most studies conclude that precipitation is indirectly correlated with amphistomy because drier habitats also tend to be more open [19, 21]. Nevertheless, the fact that amphistomy can increase water-use efficiency [11, 72] suggests that it might be favoured in dry habitats, independent of other factors.

#### H4: Altitude

Anatomical surveys demonstrate that amphistomy is sometimes more common in high elevation communities compared to nearby low elevation communities [73, 74, 75], possibly because lower CO_2_ partial pressures place a greater premium on efficient diffusion. However, this hypothesis is complicated by the fact that diffusion coefficients are higher at elevation because the air is thinner [76], meaning that CO_2_ diffusion could actually be less limiting.

#### H5: Growth form

Independent of leaf anatomy and the abiotic environment, the strength of selection on photosynthetic rate might be stronger among certain growth forms (e.g. forbs vs. trees) because of their different life history strategies. Salisbury (1927) noted qualitatively that herbs tended to amphistomatous, an observation later confirmed by Peat and Fitter (1994). However, other reviews have argued that stomatal ratio is not closely connected with any particular growth form [32, 12].

Two hypotheses I have not considered because of methodological limitations are that amphistomy is associated with vertically-oriented, isobilateral leaves [32] and that amphistomy, by doubling the conductive leaf surface area, relieves a constraint the stomatal size-density tradeoff [77, 59]. I did not have sufficient, reliable information on leaf orientation and guard cell size to evaluate these hypotheses.

#### Costs of upper stomata

This study reaffirms at a global scale that most species are hypostomatous. The most parsimonious explanation for the preponderance of hypostomy is that there is cost to having stomata on the upper surface of the leaf. A fitness cost associated with increased evaporation [78] cannot explain the dearth of stomata on the upper leaf surface, though this explanation occasionally appears in the literature [79]. In fact, amphistomy is common in some dry habitats [17, 11, 19, 20] and amphistomatous plants can be functionally hypostomatous when stressed by regulating stomatal aperture differentially on each surface [80, 81, 82, 72]. Although amphistomatous plants can be functionally hypostomatous, the reverse is not true. Hence, anatomical amphistomy should be favoured whenever the capacity to be functionally amphistomatous is advantageous.

Besides evaporation, several fitness costs have been suggested, including decreased water-use efficiency of amphistomy in large leaves [11], photodamage to guard cell chloroplasts (W.K. Smith, pers. comm.), occlusion of upper stomata by water blockage [83], and increased susceptibility to foliar pathogens [13]. Increased evaporation is an unlikely explanation since so many desert species are anatomically amphistomatous (see above), but to my knowledge, most other hypotheses have not been rigorously tested. However, [23] showed that adaxial (upper) stomata pore area, but not abaxial (lower) pore area, was strongly correlated with susceptibility to a rust pathogen. Hence, the pathogen susceptibility hypothesis is best supported by the current data.

### Text S2: Data Sources

1. Boeger and Gluzezak 2006 [84]
2. Brodribb *et al*. 2013 [59]
3. Camargo and Marenco 2011 [85]
4. Cooper and Cass 2003 [86]; Cooper *et al*. 2004 [87]
5. Dickie and Gasson 1999 [88]
6. Dunbar-Co *et al*. 2009 [89]
7. Fahmy 1997 [90]
8. Fahmy *et al*. 2007 [91]
9. Fontenelle *et al*. 1994 [92]
10. Giuliani *et al*. 2013 [60]
11. Holbrook and Putz 1996 [93]
12. Körner *et al*. 1989 [75]
13. Lohr 1919 [71]
14. Loranger and Shipley 2010 [94]
15. Malaisse and Colonval-Elenkov 1982 [95]
16. Maricle *et al*. 2009 [31]
17. Muir *et al*. 2014 [44]
18. Parkin and Pearson 1903 [96]
19. Peace and MacDonald 1981 [97]
20. Rao and Tan 1980 [98]
21. Reed *et al*. 2000 [99]
22. Ridge *et al*. 1984 [100]
23. Selvi and Bigazzi 2001 [101]
24. Seshavatharam and Srivalli 1989 [102]
25. Sobrado and Medina 1980 [103]

### Text S3: An evolutionary process model for proportion traits

Making evolutionary sense of a biological pattern requires an underlying process model to provide the theoretical foundation on which data analysis rests. A powerful approach in macroevolution involves modelling trait evolution on adaptive landscapes where the peaks of high fitness evolve with or without constraint [28, 104, 105]. If models with constraint describe the data better than those without, then there is compelling evidence that the adaptive landscape is shaped by some combination of selective, genetic, functional, or developmental constraints. Furthermore, the adaptive landscape may change under multiple selective regimes, meaning that a trait is best described by a mixture of distributions, each generated under separate selective regimes [27, 29, 30]. Current evolutionary process models such as Brownian motion and Ornstein-Uhlenbeck assume that traits follow a Gaussian distribution, but this is clearly inappropriate for traits like stomatal ratio. In this text, I modify previous evolutionary process models to accommodate proportion traits and derive the expected pattern given adaptive landscapes that are constrained versus those that are unconstrained. This model provides a strong theoretical foundation for the model-based statistical inference described in Text S4. A glossary of symbols used in this text are provided in Table S1.

**Table S1.** Glossary of symbols used in process models of stomatal trait evolution.

| Symbol | Description |
| --- | --- |
| $r$ | Stomatal ratio: ratio of upper to total stomatal density |
| $N_T, N_U, N_L$ | Number of stomata in a focal leaf area $A_L$<br>The total number $N_T$ is the sum of upper $N_U$ and lower $N_L$ stomata |
| $D_T, D_U, D_L$ | Density of stomata in total, upper, and lower surfaces |
| $A_{\text{leaf}}$ | Focal leaf area |
| $\nu$ | Diffusion coefficient of stomatal ratio |
| $\theta$ | Long-run average stomatal ratio |
| $\alpha$ | Return rate to long-run average ratio |
| $\phi$ | Defined as $\nu\alpha$ |
| $M_{\delta x}$ | Drift function of stomatal ratio $r$ in diffusion approximation |
| $V_{\delta x}$ | Diffusion function of stomatal ratio $r$ in diffusion approximation |

In both models with and without constraint, I assume that *total* stomatal density follows a random walk over macroevolutionary time, though the exact process is irrelevant here. Imagine for a set area (*A*_leaf_) of leaf (e.g. 1 *µ*m^2^) there are *N_T_*(*t*) = *A*_leaf_*D_T_*(*t*) = *A*_leaf_(*D_U_*(*t*) + *D_L_*(*t*)), where *N_T_*(*t*) is the total number of stomata in that area at time *t*. Total stomatal number *N_T_*(*t*) is the sum of upper (*N_U_*(*t*)) and lower (*N_L_*(*t*)) stomata. Let Δ*N_T,t_* = *N_T_*(*t* + 1) − *N_T_*(*t*) be the change in total stomatal number that must be made up of changes in upper stomata, lower stomata, or some combination of both. I assume that the contribution to Δ*N_T,t_* from upper and lower stomata is proportional to their density. For reasons explained below, I define *ν* = *N_T_*(*t* + 1) as the total stomata at time *t* + 1. The transition rate *u_ij_* from *N_U_* = *i* upper stomata at time *t* to *N_U_* = *j* upper stomata at time *t* + 1 is binomially distributed with a rate determined by the stomatal ratio *r*:

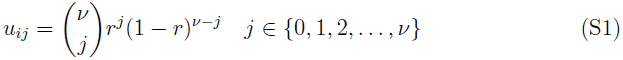

Note that stomatal ratio here is defined as the proportion of upper stomata, *r* = *N_U_*/(*N_U_* + *N_L_*) = *N_U_*/*N_T_* = *N_U_/ν*. The mean and variance of stomatal ratio in the next time step is therefore:

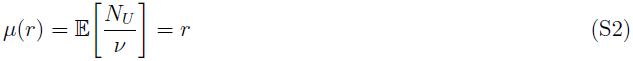

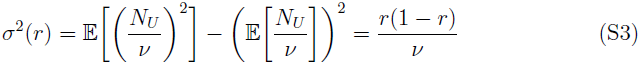

In other words, the average stomatal ratio does not change, but the variance increases each time step. When *ν* is large, the distribution can be approximated with a normal distribution and a diffusion approximation can be used to model the long term evolution of the trait. This diffusion process is analogous to Brownian motion, except that the trait is bounded by 0 and 1. It is also mathematically equivalent to one-locus, two-allele population genetic models of neutral evolution (see [106] for a detailed derivation). I will make reference to results from this literature without rigorously deriving them here. In particular, it has been shown that the stationary distribution of the diffusion is:

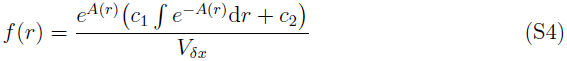

where

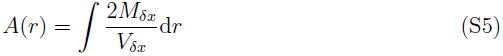

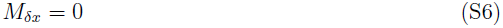

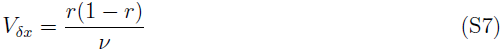

and the time scale is in units of *ν*^−1^. Thus, *ν* can be interpreted as a diffusion coefficient without necessarily specifying a genetic or developmental mechanism that governs the amount of variance in stomatal ratio from one time to the next. Solving for *f*(*r*) without selection on stomatal ratio yields:

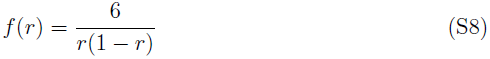

Thus, without selection on stomatal ratio, most species should be hypo- or hyperstomatous (Fig. S3). Next, I modify the model to include stabilizing selection around a long-run average *θ*, which may be interpreted as a peak in the adaptive landscape under a single selective regime. This process model is analogous to an Ornstein-Uhlenbeck process for a bounded trait. I again use the diffusion approximation, but this time the drift and diffusion coefficients are:

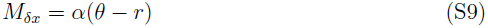

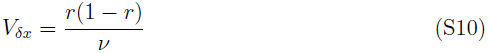

*α* is the return rate to *θ*. Greater values of *α* constrain trait variation more tightly around *θ*. With these coefficients and setting the first constant of integration *c*_1_ to 0 yields:

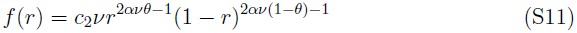

where:

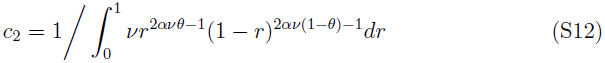

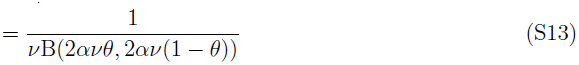

B(·) is the beta function. Setting *c*_1_ to 0 can be justified by recognizing that the distribution should be symmetrical (*x* = 1 − *x*) when *θ* = 0.5, which only occurs if *c*_1_ = 0 (S.P. Otto pers. comm.). Further, I confirmed the accuracy of the analytically-derived stationary distribution using stochastic simulations (data not shown).

Defining *φ* = *αν*, the stationary distribution simplifies somewhat to:

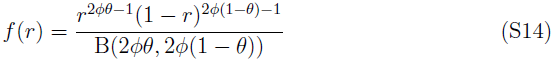

This is the Beta(*α*, *β*) distribution with *α* = 2*φθ* and *β* = 2*φ*(1 − *θ*). Note that, following standard notation, *α* here refers to the first shape parameter of the Beta distribution, not the constraint factor of the evolutionary process model. This result means that the well-known statistical properties of the Beta distribution can be leveraged to understand the stationary distribution of a proportion trait under a constrained adaptive landscape. For example, the Beta distribution takes on a variety of shapes that begin to resemble the distribution of proportional traits like stomatal ratio (Fig. S4). Hence, the evolutionary process model developed here provides a strong theoretical justification for fitting the stomatal ratio data to a mixture of Beta distributions in order to infer the selective regimes shaping this trait across plant species. Although I have derived the model with stomatal ratio in mind, it should be applicable to wide variety of proportional traits evolving under a constrained adaptive landscape.

**Fig. S3.**
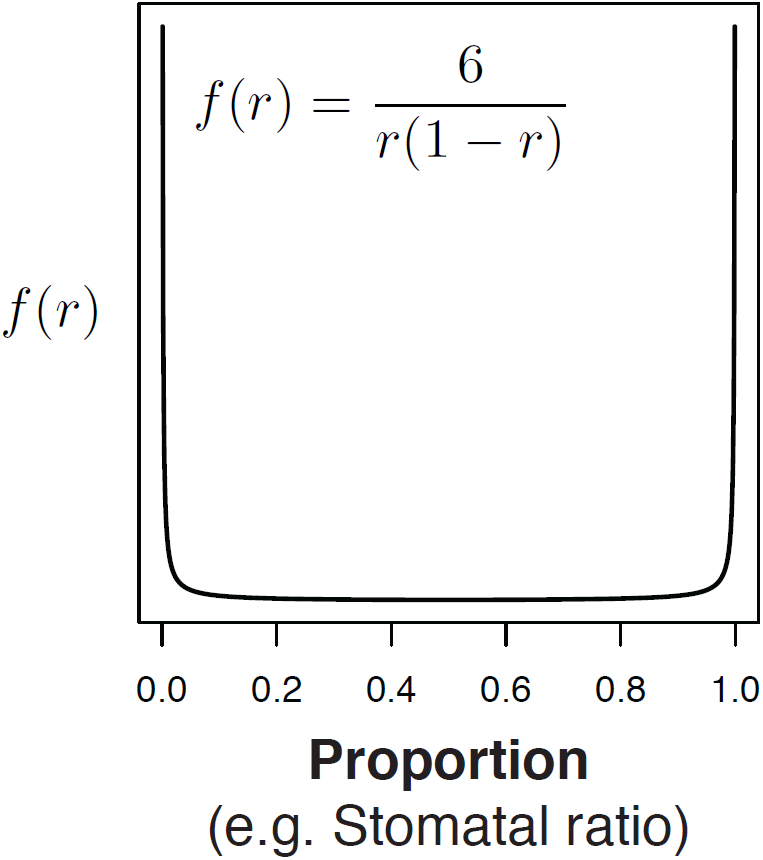
Without constraint, a proportion trait like stomatal ratio (*r*) will evolve toward a distribution in which most species are 0 or 1.

**Fig. S4.**
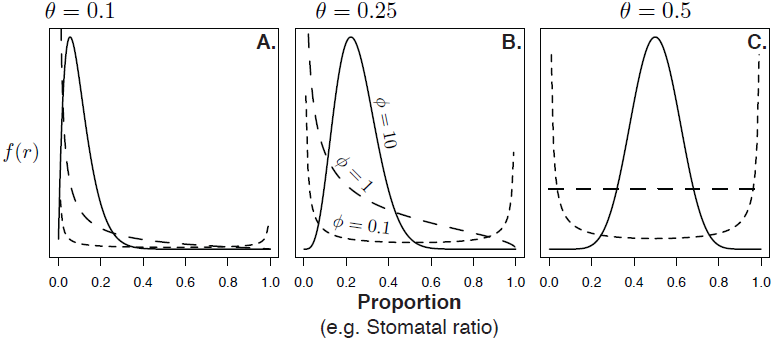
A proportion trait like stomatal ratio evolving under a constrained adaptive landscape is Beta distributed. The Beta distribution can take on a wide variety of shapes depends on the long-run average *θ* and the levels of constraint *φ* (greater *φ* equals greater constraint).

### Text S4: Fitting evolutionary process to pattern using finite mixture models estimated with maximum likelihood

In this paper, I infer the number of selective regimes acting on stomatal ratio by fitting a mixture of stationary distributions derived from the process model above to the data. In this section I derive the likelihood functions and describe an expectation-maximization algorithm to find the maximum likelihood mixture model given the data. R code to implement these methods is available on Dryad [62]. In general, finite mixture distributions are the summation of *k* ≥ 2 mixture components (i.e. probability distributions) with density *f_i_*(*x*) and mixture weight *w_i_*:

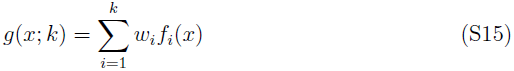

Here the *i*-th mixture component has a probability density *f_i_*(*x*) given by the stationary distribution in Eq S14 with parameters *θ_i_*, *φ*_*i*_. The likelihood of a mixture distribution given *k* mixture components and a data vector ***x*** with sample size *n* is the weighted sum of the likelihoods of each component:

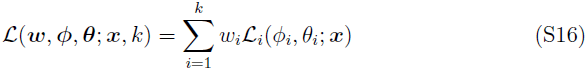

The parameter vectors ***w***, *φ*, and ***θ*** are defined as:

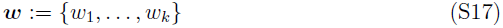

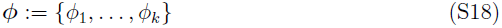

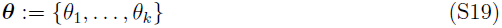

For the *i*-th component, the likelihood of parameters *φ*_*i*_ and *θ_i_* given the data is the product of the probability densities of each datum (*x*_1_, *x*_2_, …, *x_n_*):

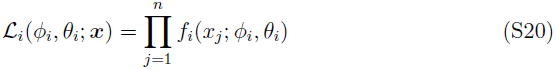

To obtain reasonable fits, I found it necessary to modify the likelihood to incorporate left- and right-censored data. This is because the stomatal ratio dataset contains many 0’s (all stomata are on the lower surface of the leaf) and 1’s (all stomata on the upper surface). Under most parameterizations of the Beta distribution, the probability density of 0 and 1 is ∞ or 0. I left- and right-censored the data at *x*_1_ = 0.001 and *x_r_* = 0.999 as these were very close to the lowest and highest values reported in the dataset (except 0 and 1), respectively. This means that any datum reported as 0 was statistically interpreted as falling anywhere between 0 and 0.001. Likewise, a datum reported as 1 was assumed to fall between 0.999 and 1. A reasonable interpretation is that a stomatal ratio so close to 0 or 1 would be practically difficult to measure. Biologically, a stomatal ratio less than 0.001 or greater than 0.999 are indistinguishable from 0 and 1. With censoring, the likelihood of the *i*-th component becomes:

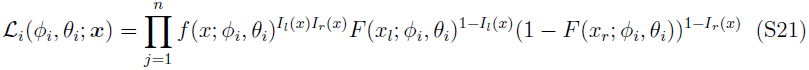

*F*(*x*; *φ*_*i*_, *θ_i_*) is the cumulative density function of the Beta distribution; *I_l_*(*x*) and *I_r_*(*x*) are indicator functions:

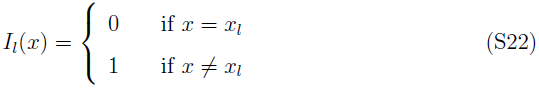

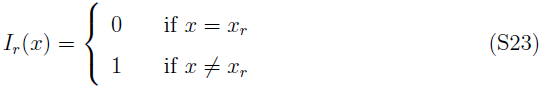

To find the maximum likelihood mixture distribution, I used an expectation-maximization (EM) algorithm similar to [107]. EM algorithms are particularly well-suited to fitting mixture distributions. Here, I describe the initialization, expectation (E-step), and maximization (M-step) procedure.

#### Initialization

The data were divided into *k* evenly-sized components. For example, if *k* = 2, data below the median were assigned to component 1; data above the median were assigned to component 2. For each component, the initial weight was therefore *w*_*i*,init_ = 1/*k*. Within each component, I used the optim function in R to estimate the maximum likelihood parameters 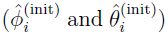 of a Beta distribution. Note that I am using parenthetical superscript to indicate the iteration of the algorithm, starting with the initial parameterization, followed by t = 1,2,3,… until the likelihood converges. The initial parameter vectors are therefore:

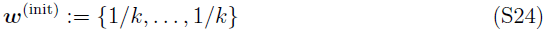

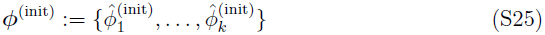

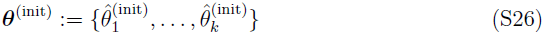

#### Expectation

In the E-step, the expected likelihood is calculated under the parameters estimated from the previous iteration. The mixture weights are then updated and carried forward to the M-step. For the first iteration following initialization, the mixture weights ***w***^(1)^ conditional on the initial parameterization are:

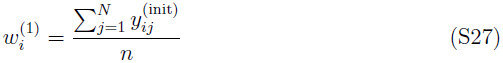

where 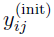 is the probability that *x_j_* belongs to component *i* given initial parameters:

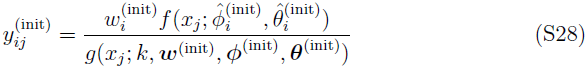

In subsequent iterations, the equations are similarly:

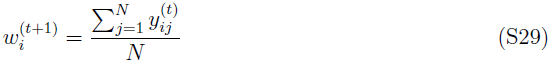

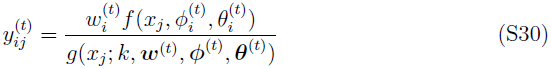

#### Maximization

During the M-step, estimates of *φ* and ***θ*** are updated using maximum likelihood conditional on mixture weights calculated in the E-step:

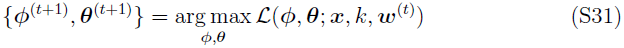

I used the optim function in R to find the parameters that maximized the likelihood function. After the M-step, the next iteration begins at the E-step and continues until the likelihood converges to a stable value. As with other hill-climbing likelihood searches, EM does not guarantee convergence at the maximum likelihood. With the stomatal ratio data, I found that multiple initialization procedures yielded the same final parameter estimates, suggesting that the algorithm was successfully converging on the maximum likelihood solution.

